# Revealing pan-cancer clonal niches of T cells from single-cell RNA sequencing using contrastive learning

**DOI:** 10.64898/2026.10.06.757206

**Authors:** Ofir Shorer, Asaf Pinhasi, Ron Amit, Keren Yizhak

## Abstract

T-cell clonal organization is central to anti-tumor immunity, yet its functional characterization typically requires paired single-cell RNA and TCR sequencing (scRNA/TCR-seq). Existing multimodal methods integrate transcriptome and TCR information but are limited when TCR data are unavailable. Here we present CloneTrast, a supervised contrastive-learning framework that uses paired scRNA/TCR-seq during training to learn clonal-functional states from gene expression, while requiring only scRNA-seq at inference. Using paired scRNA/TCR-seq profiles from 2.6 million human T cells across 16 cancer types, CloneTrast identified distinct T-cell clonal niches not explained solely by TCR sequence similarity or conventional cell-state annotations. Among these, an NK-like T-cell niche enriched for tumor-blood shared TCRs was consistently associated with poor immunotherapy response at the pan-cancer level. Analysis of antigen-specific T-cell clones revealed that tumor-resident and blood-borne antigen-specific T cells were preferentially associated with distinct clonal niches. Overall, CloneTrast enables scalable and interpretable inference of clonal organization from single-cell transcriptomes.

## Introduction

T cells play a central role in anti-tumor immunity, where their functional states, clonal expansion, and antigen specificity, collectively shape immune recognition and response to therapy^1–4^. Single-cell RNA sequencing (scRNA-seq) has enabled high-resolution characterization of T-cell phenotypes, revealing diverse differentiation states across tumor and blood samples with direct clinical relevance^5–10^. In parallel, single-cell T-cell receptor sequencing (scTCR-seq) provides clonal information by linking individual T cells to their receptor sequences, allowing the identification of expanded clones, tracking of clonal dynamics, and inference of relationships between tumor-infiltrating and circulating T-cell populations^11^. Together, paired scRNA/TCR-seq has substantially improved our ability to study how T-cell clones differentiate, expand, migrate, and respond to cancer therapy^12–15^.

Several studies have shown that T-cell clonal dynamics are associated with immunotherapy response across cancer types^14–20^. Characteristics such as T-cell abundance, expansion, replacement, reinvigoration, and phenotypic states of tumor-infiltrating T-cell clones have all been linked to response or resistance to immune checkpoint blockade^12,13,15^. These findings highlight the importance of studying T-cell function not only at the level of individual transcriptional states, but also at the level of clonally related T cells. Yet clonal identity and transcriptional states are not uniquely coupled: T cells within the same clone can exhibit substantial phenotypic heterogeneity, while transcriptionally similar T cells can arise from many different clones^21,22^. Therefore, a major challenge is to define functional relationships between T-cell clones in a way that integrates clonal organization with transcriptional states.

To address this challenge, multiple computational methods have been developed for the joint analysis of paired scRNA/TCR-seq data. These approaches have demonstrated that combining gene expression and TCR information can provide a richer representation of T-cell biology than either modality alone^23–29^. Some methods learn shared low-dimensional representations of single-cell transcriptomes and TCR sequences^23–27^, whereas others operate at the clonotype-level or use graph-based strategies to relate transcriptional and receptor similarity^28,29^. Collectively, these tools have advanced the analysis of antigen-specific T-cell populations, clonotype-associated transcriptional programs, and multimodal T-cell states.

Despite these advances, important limitations remain. First, most existing approaches require paired gene expression and TCR information for each cell during inference. This requirement restricts their use in datasets with low or incomplete TCR capture, as well as across large public scRNA-seq resources where only transcriptomic profiles are available. Second, many methods learn dataset-specific representations and therefore require retraining or re-optimization for each new dataset, limiting direct cross-study applications. Third, approaches that rely directly on TCR sequence similarity may be less suited to identifying convergent transcriptional programs shared by T-cell clones with distinct receptor sequences. Finally, the limited interpretability of many representation-learning models makes it difficult to identify the transcriptional features that define the learned clonal-functional structure.

Here, we developed CloneTrast, a supervised contrastive-learning framework designed to learn a transcriptome-based representation of T-cell clonal function. CloneTrast is trained on paired scRNA/TCR-seq data, using TCR information to define clonal identities during training. TCR sequences are used solely to provide clonal labels for supervision and are not included as model inputs. Instead, the model learns a contrastive representation of gene expression in which cells belonging to the same clone are mapped closer together, while cells from different clones are pushed apart. Once trained, CloneTrast can be applied to scRNA-seq data alone, enabling the inference of clonal relationships from transcriptional profiles in datasets lacking paired TCR sequencing. We trained and evaluated CloneTrast using our in-house-constructed human pan-cancer resource of paired scRNA/TCR-seq profiles, including more than 2.6 million T cells from tumor and blood samples across 16 cancer types. CloneTrast identified groups of T-cell clones with shared functional properties, defined as **clonal niches**. These niches capture an intermediate level of organization between individual clonotypes and conventional T-cell state annotations: they are neither defined solely by identical or sequence-similar TCRs, nor fully explained by standard transcriptomic annotations. Using model explainability, we characterized the transcriptional programs underlying niche identity and identified niche-specific features that extend beyond standard differential gene expression.

Applying CloneTrast to both training and external test datasets, we uncovered two patterns of clonal-functional organization spanning tumor and blood samples: (1) clonal niches composed primarily of a single major T-cell differentiation state, termed ‘identity niches’, and (2) clonal niches composed of multiple T-cell differentiation states altogether, termed ‘functional niches’, showing intriguing non-conventional gene expression patterns. We further found that specific clonal niches are associated with patient response to immunotherapy at the pan-cancer level. Finally, in HBV-positive hepatocellular carcinoma patients^30^, CloneTrast identified a clonal niche enriched for high-avidity HBV-specific T-cell clones, suggesting that clonal niches can capture biologically meaningful antigen-related T-cell programs. Analysis of antigen-specific T cells across additional patient cohorts^31–34^, spanning different cancer types and antigens, revealed that tumor-resident and blood-borne antigen-specific T cells are preferentially associated with distinct clonal niches. Together, CloneTrast provides a scalable and interpretable framework for studying T-cell clonal functional organization from single-cell transcriptomes.

## Results

### CloneTrast dissects the functional states of T-cell clones using single-cell gene expression

To train a model that captures the clonal functional states of T cells at the pan-cancer level, we first collected publicly available data of 1,451 tumor and blood samples obtained from cancer patients across 16 cancer types, consisting of 37 datasets with more than 2.6 million T cells passing a strict quality control done separately on both scRNA-seq and scTCR-seq data^12,13,30,31,35–65^ (Table S1, Method details). Overall, 26 datasets containing 963 samples across 14 cancer types, with more than 1.5 million T cells, were used to train the model (Figure 1A), with an additional 1 million T cells across 488 samples from 11 other datasets, including two cancer types unseen during training, were used to test model performance (Table S1). Across the train set, 11,950 genes shared between all datasets were used for further integration of all datasets (Figure S1A-E, Method details). Single cells were labeled according to their clonal identity based on CDR3 sequences, and their clone size was quantified accordingly (Method details).

**Figure 1.**
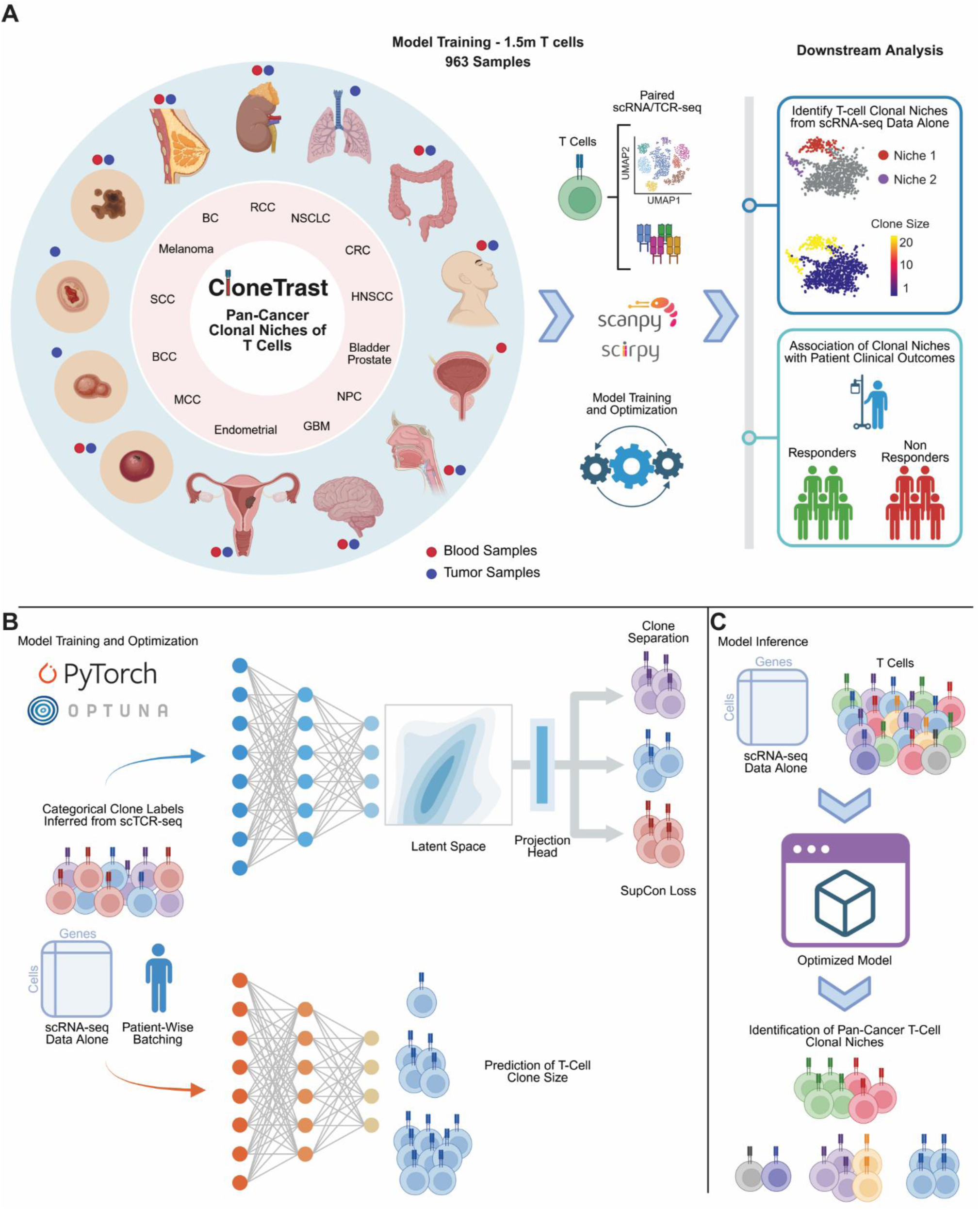
CloneTrast dissects the functional states of T-cell clones using single-cell gene expression. A. A schematic workflow of the study depicting T-cell data collection across tumor and blood samples of human cancer patients, strict quality control, model training and optimization, as well as downstream analysis, including identification of pan-cancer T-cell clonal niches and their association with patient clinical outcomes. B. A schematic description of the model architecture, which learns a low-dimensional representation of T-cell clones using supervised contrastive loss, with a complementary model for prediction of T-cell clone size. The contrastive model was trained using patient-wise batching to improve both inter- and intra-patient performance, ensuring the model learns meaningful features within individual patients. C. A schematic description of model inference using scRNA-seq data alone. See also Table S1.

CloneTrast was designed to use gene expression alone in order to learn a latent representation that maximizes the cosine similarity of T cells, based on their clonal identity, using supervised contrastive loss^66^ (Figure 1B, Method details). Specifically, TCR data are used during training to define clonal relationships: cells from the same clone serve as positives, cells from different clones serve as negatives, and the model learns from gene expression alone. This specific modeling approach results in a distinct embedding space capturing multiple components of clonal organization (Figure 1C): **(1) Different clones, similar functional states:** Two clones defined by different TCR sequences can be positioned near one another if they share similar functional status; **(2) Same clone, divergent states:** T cells belonging to the same clone are pulled toward one another despite having diverse differentiation states due to intra-clonal transcriptional heterogeneity; **(3) Same TCR sequence, different functional states:** Two clones with identical TCR sequences but distinct clonal identities, because they originate from different patients, can be positioned far apart if they exhibit different functional states.

Simultaneously, we trained an additional complementary model for prediction of T-cell clone size using gene expression alone, providing an additional layer of information upon the learned representation of clonal niches (Figure 1B, Method details). Both the contrastive and the clone-size prediction models are fully integrated into the CloneTrast package, and can be used immediately for downstream data analysis.

### Learning pan-cancer low-dimensional representation of T-cell clones using CloneTrast

In order to train a large-scale model at the pan-cancer level, we used our training set of 1.5 million T cells and allocated 80% of the data for training and 20% for validation. The validation set was used to evaluate model performance for hyperparameter tuning using Optuna^67^ (Method details). Of note, cell allocation was done while enforcing that all the cells from a given patient are assigned exclusively to either training or validation set, but not both, thus preventing data leakage (Method details). Model optimization was performed while aiming to maximize the weighted per-patient clonal purity score across all training trials (Figure S2A, Method details). This optimization score is designed such that for each patient, it quantifies how well T cells from the same clone are pulled together, by measuring for each cell the fraction of its 10 nearest neighbors having the same clonal identity upon the learned representation (Method details). Thus, besides minimizing the supervised contrastive loss based on clonal identity of T cells, we also maximize the per-patient grouping of T-cell clones by adjusting the model hyperparameters (Figure S2A, Method details). Following, the best optimized model, based on validation-set performance, was used to obtain a learned representation of the data using the entire training set of 1.5 million T cells (Figure 2A). Importantly, this new low-dimensional representation of T cells is different from the representation obtained using expression-based workflows for processing scRNA-seq data alone, as it explicitly aims to bring together T cells sharing the same clonal identity based on their gene expression profile (Figure 2B). Accordingly, CloneTrast substantially improved the patient clonal purity score compared with the standard representation of an expression-based embedding (p-value = 2.24·10^-54^, Figure 2C, Method details). Complementing this representation-learning framework, we applied the trained and optimized model for prediction of T-cell clone size from gene expression (Method details). The predicted clone size showed a significant correlation with the observed clone size across T-cell clones (Spearman-r = 0.799, p < 2.23·10^-308^, Figure 2D, Method details).

**Figure 2.**
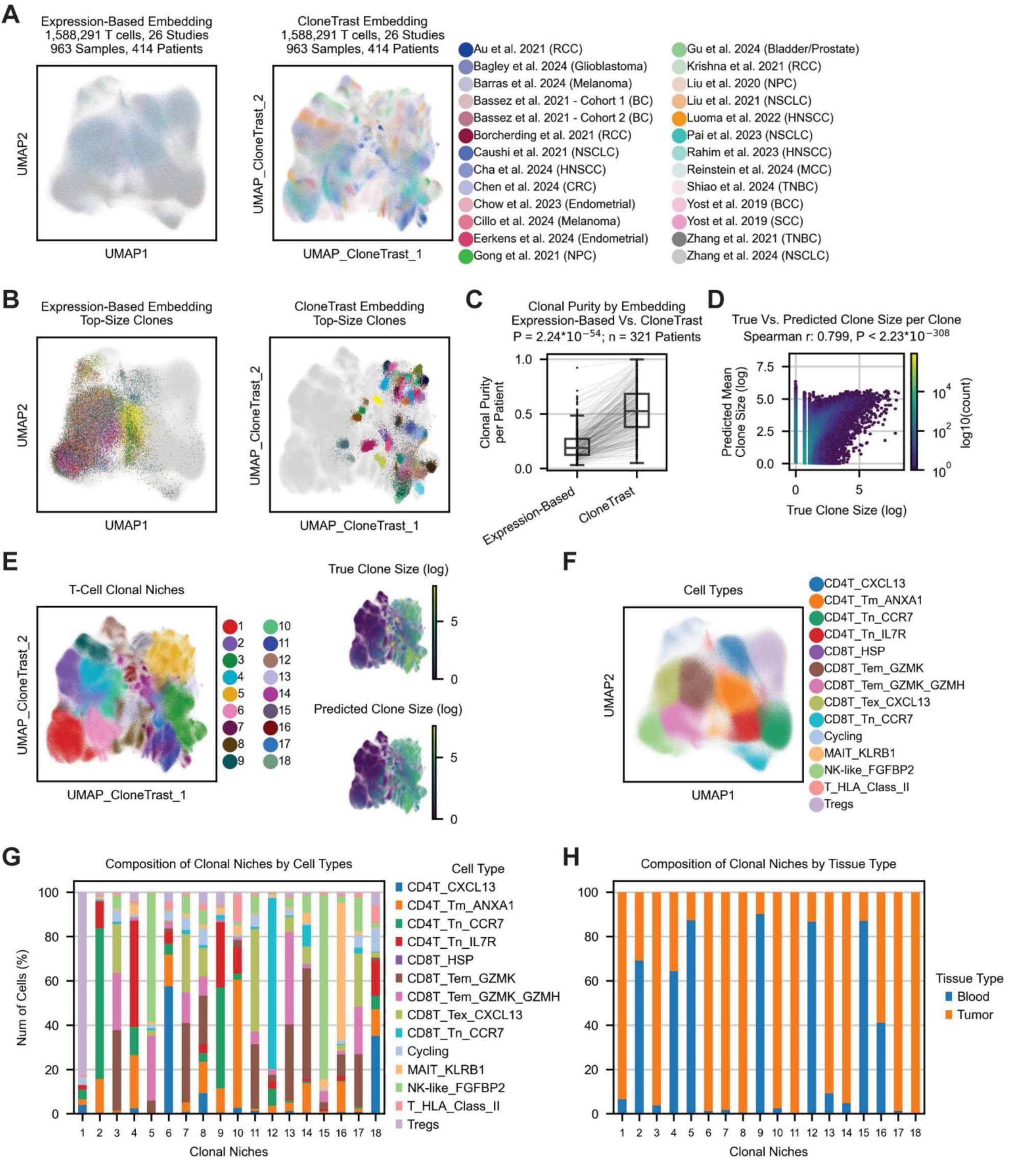
Learning pan-cancer low-dimensional representation of T-cell clones using CloneTrast. A. Low-dimensional representation of the entire training set, composed of more than 1.5 million T cells, colored by 26 different studies. Cells are presented using an expression-based representation obtained using a standard processing pipeline for gene expression data (left), as well as by the representation obtained using CloneTrast (right). B. Expression-based representation (left) and CloneTrast-based representation (right), showing the exact same T cells from top-size clones having clone size larger than 500 T cells. C. Paired boxplots demonstrating the difference of clonal purity score per patient, calculated using either an expression-based embedding or the embedding obtained using CloneTrast (p-value = 2.24·10^-54^, n = 321, paired two-sided Wilcoxon signed-rank test). Only patients having clones with a minimal size of 20 cells were used for this calculation. D. Scatter plot showing the Spearman correlation between the observed clone sizes and those predicted by our model (Spearman-r = 0.799, p-value < 2.23·10^-308^). E. Leiden clustering results representing 18 distinct T-cell clonal niches upon the learned representation of CloneTrast (left), including overlay of both the observed and the predicted clone size for each T cell (right). F. Expression-based representation of T cells labeled by different T-cell states/types. G. Stacked barplot showing the composition of each clonal niche by different T-cell types. H. Stacked barplot showing the composition of each clonal niche by tissue type of origin, including tumor and blood samples. See also Figures S1-S2 and Tables S1-2.

To characterize the phenotypic landscape of T cells captured by CloneTrast in an unbiased manner, we clustered T cells within the learned representation using the Leiden algorithm^68^ (Figure 2E, Method details). This step resulted in 18 distinct clusters of T-cell clones having varying levels of clone sizes, as well as different composition of T-cell states, spanning both tumor and blood samples (Figure 2E-H, Table S2). These clusters represent distinct functional states of T cells at the clonal level, which we term ‘clonal niches’. As such, rather than reflecting individual T-cell differentiation states, these clonal niches capture shared transcriptional programs of T-cell clones, revealing different patterns of clonal functional states. For example, the population of CXCL13-expressing CD8^+^ T cells, which was previously linked to immunotherapy outcomes across multiple cancer types^14,69^, spans multiple different clonal niches, including niches 3, 7, 8, 11, 13, 14, and 17 (Figure 2E-H), reflecting that the same differentiation state can be considered in different clonal contexts. Similarly, differentiation states such as GZMK- and GZMH-expressing effector-memory CD8^+^ T cells, NK-like T cells, and cycling T cells, span multiple different clonal niches (Figure 2E-H). In contrast, differentiation states such as Tregs, naive CD4^+^ T cells, naive CD8^+^ T cells, and CXCL13-expressing CD4^+^ T cells, show more restricted localization, typically occupying only one or two niches (Figure 2E-H). These results indicate that conventional differentiation state annotations do not fully capture the functional organization of T-cell clones. Instead, the clonal niches captured by CloneTrast provide an orthogonal layer of biological organization, revealing distinct clonal contexts within shared differentiation states.

Because CloneTrast is trained using clonal identities without feeding the actual TCR sequence to the model, we next asked whether the resulting clonal niches may reflect similarity in TCR sequences, or instead, capture broader transcriptional states shared across distinct T-cell clones. To address this, we applied GLIPH2^70,71^ for grouping TCRs based on CDR3 sequence similarity across datasets, rather than relying on exact CDR3 sequence identity (Figure S2B). As expected, we found that CloneTrast captures convergent transcriptional phenotypes of T-cell clones that may arise from different TCR sequences. Consistent with this interpretation, although clonal diversity within clonal niches was significantly lower when calculated using GLIPH2-defined CDR3 groups compared to the original clonal identities (p-value = 7.63·10^-6^, Figure S2C), the overall diversity within each niche remained high (normalized Shannon entropy = 0.756-0.998, Table S2). These findings support the notion that CloneTrast groups T-cell clones based on shared functional status, largely independent of CDR3 sequence identity.

In order to demonstrate the distinction of our approach from existing representation-learning methods of paired scRNA/TCR-seq data, we applied mvTCR^23^ and MIST^25^ using our training set of 1.5 million T cells (Method details). These two methods were the only scalable methods that could process this amount of T cells in reasonable run times. Importantly, the multimodal models implemented by both methods require paired scRNA/TCR-seq data during model inference to generate a joint representation of the two modalities, a scenario distinct from CloneTrast’s ability to infer clonal-functional structure from scRNA-seq data alone. While the low-dimensional representations obtained using mvTCR and MIST preserved the overall clonal structure of individual T-cell clones (Figure S2D), these representations resulted with clones attributed to the same functional state in different locations, as well as clones having different functional states in the same location (Figure S2E-G). In addition, CloneTrast maintained effective batch integration across samples, with performance approaching that of MIST and mvTCR (Figure S2H, Table S2, Method details), despite learning a representation explicitly optimized to organize T cells according to clonal identity. This indicates that the clonal structure captured by CloneTrast is not driven primarily by sample-specific effects. Another important distinction lies in the gene space used by these models. MIST and mvTCR are trained using the top 2,000 and 5,000 highly variable genes, respectively, selected specifically from the training dataset, whereas CloneTrast uses a fixed set of 11,950 quality-controlled genes. At inference, CloneTrast automatically aligns new datasets to this fixed gene space by matching shared genes and accounting for genes absent from the query dataset (Method details). This enables direct application of the pre-trained model across independent studies without redefining the input feature space or retraining.

Taken together, these results demonstrate that CloneTrast learns a robust, clonally informed expression-based representation of T-cell functional states that is distinct from both conventional transcriptomic representations and existing multimodal approaches.

### Characterization of T-cell clonal niches at the pan-cancer level using model interpretability

To characterize the functional status of each clonal niche, particularly given that individual niches can comprise multiple different T-cell states altogether (Figure 2G), we used Captum^72^, a model interpretability library for PyTorch^73^ enabling memory-efficient and scalable computations (Figure 3A, Method details). We applied Captum to identify the features that most strongly define each niche by promoting similarity among T cells within the niche, while distinguishing them from cells in other niches. This approach goes beyond standard differential gene expression and can reveal complex, non-linear patterns affecting model behavior. Genes may be differentially expressed between T-cell populations without necessarily being the features that determine the clonal organization of these populations. Conversely, genes having context-dependent effect on the functional clonal organization of T cells, may contribute substantially to the separation of clonal niches without being the most significantly differentially expressed genes in those niches. This approach identified a ranked set of genes that define each niche, capturing gene expression profiles underlying niche identity (Table S2).

**Figure 3.**
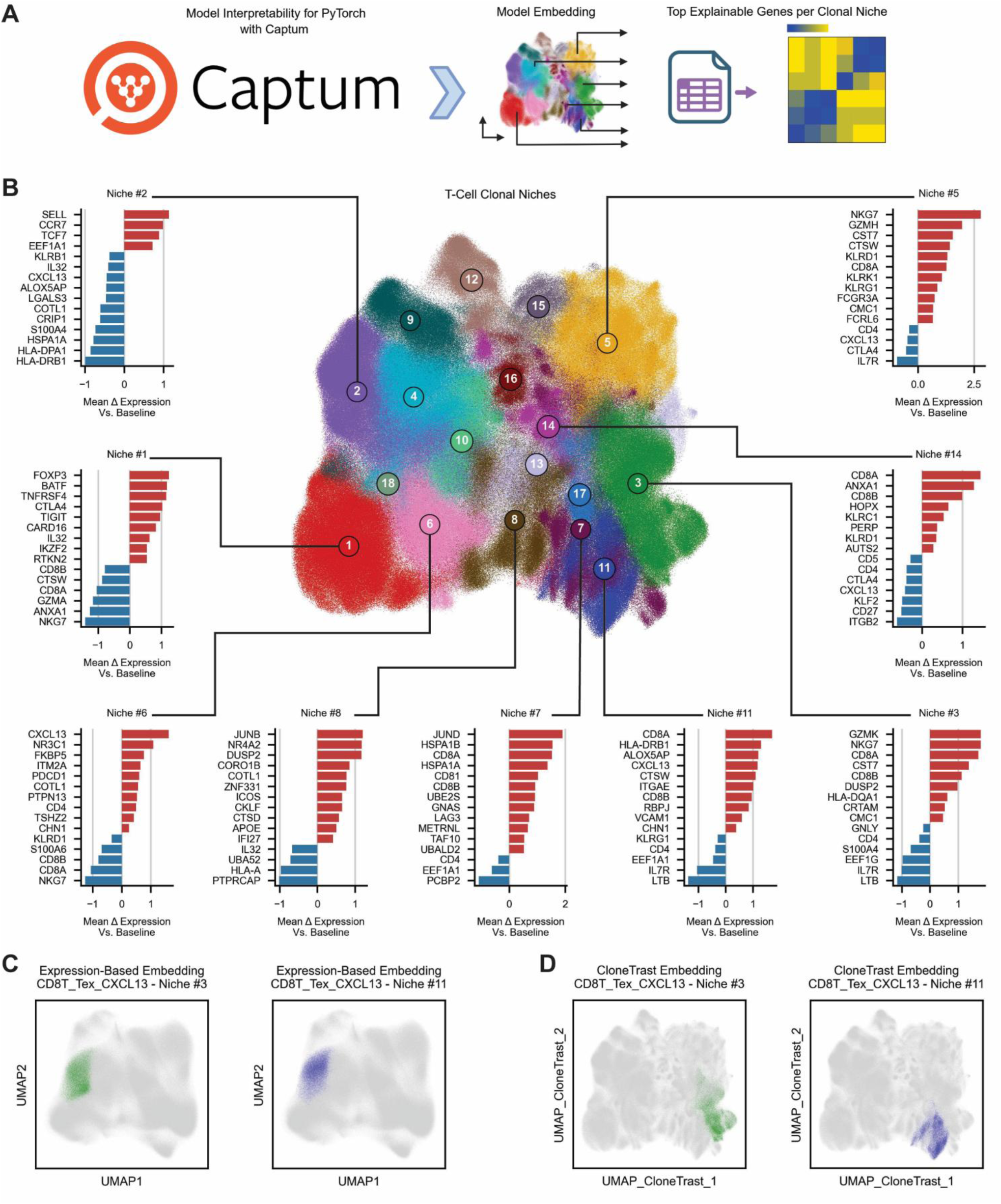
Characterization of T-cell clonal niches at the pan-cancer level using model interpretability. A. A schematic workflow depicting the application of Captum^72^ to obtain the most explainable features driving T-cell clonal niche identity. B. Representation of the top 15 explainable genes across selected clonal niches, ranked within each niche according to the difference between their mean expression in the niche and across all cells in the dataset. C-D. Representation of CXCL13^+^ CD8^+^ T cells from selected clonal niches, including niches #3 and #11, respectively, upon the expression-based representation obtained using the common processing pipeline for gene expression data (C), and upon the representation obtained using CloneTrast (D). See also Figures S3-S5 and Table S2.

We first identified clonal niches that are predominantly composed of a single canonical T-cell population, which we termed ‘identity niches’. These include niche #1, composed mostly of Tregs, niche #2 of naive CD4^+^ T cells, niche #12 of naive CD8^+^ T cells, niche #15 of NK-like T cells, and niche #16, composed primarily of mucosal-associated invariant T (MAIT) cells (Figure 2G). Consistent with their well-defined cellular identities, their top explainable genes closely resemble the set of their top differentially expressed genes (Figure 3B, Table S2).

In contrast, other clonal niches were composed of multiple T-cell differentiation states, which we termed ‘functional niches’, demonstrating intriguing non-conventional gene expression patterns. A notable example includes CXCL13^+^ CD8^+^ T cells that are clustered together upon the expression-based representation (Figures 3C, S3A). However, CloneTrast divides this population between five different niches, #3, #7, #8, #11 and #17, thus capturing five different sub-populations of CXCL13^+^ CD8^+^ T cells differed by their clonal functional status (Figure 3D, S3B-C). Notably, expression of *CXCL13* is found to be highly explainable only for niche #11 despite being differentially and highly expressed in all five niches (Table S2). The top explainable genes of each clonal niche further capture different functional states of these sub-populations: the genes underlying niche #3 identity include *NKG7*, *GZMK*, *CST7*, *CMC1* and *DUSP2*, consistent with a more effector-memory phenotype; *ITGAE* and *ZNF683* which underlie niche #11 identity, mark a tissue-resident phenotype; in addition, niche #7 is characterized by the expression of multiple heat-shock protein (HSP) genes, indicating a subpopulation of CXCL13^+^ CD8^+^ T cells exhibiting a stress response phenotype; the genes *IFI27*, *IFITM1*, *IFITM2*, *IFI30*, and *IFI44L*, which indicate interferon-response program, together with *JUNB* and *ICOS*, which are associated with TCR activation, underlie niche #8 identity; lastly, the genes *KLRK1*, *KLRC4*, *KLRC2*, *TNFRSF9*, *PRF1*, and *NKG7*, all associated with T-cell cytotoxicity, together with *CXCR4* and *ITGA4*, which are involved in T-cell trafficking, underlie niche #17 identity (Table S2).

Importantly, these distinctions were not simply a consequence of differences that were readily apparent in standard transcriptional space. When we restricted the analysis exclusively to CXCL13^+^ CD8^+^ T cells and repeated a conventional expression-based dimensionality reduction, the cells formed a largely continuous population rather than resolving into the distinct sub-populations identified by CloneTrast (Figure S3D-E). This was the case even though several genes contributing to the identities of individual niches were also differentially expressed among these sub-populations (Table S2). Thus, differential expression of individual genes was insufficient to recover the clonal organization captured by CloneTrast. Instead, the joint feature representation learned by CloneTrast revealed a higher-order organization that was not captured by conventional expression-based dimensionality reduction.

The same principle extended beyond CXCL13^+^ CD8^+^ T cells. Multiple other transcriptionally defined T-cell populations were distributed across multiple functional niches, indicating that this phenomenon was not restricted to a single differentiation state. At the same time, individual niches could contain cells belonging to multiple different T-cell populations. For example, NK-like T cells were distributed across multiple clonal niches, including niches that also contained additional T-cell populations, such as CXCL13^+^ CD8^+^ T cells, with each niche exhibiting distinct transcriptional programs despite the shared NK-like canonical identity (Figure S4A-C). Similarly, GZMK-expressing effector-memory CD8^+^ T cells were distributed across multiple niches, several of which also contained additional T-cell populations, showing distinct underlying transcriptional features (Figure S5A-C). Importantly, analysis of the largest clones in our data showed that these three distinct T-cell populations, including CXCL13^+^ CD8^+^ T cells, GZMK-expressing effector-memory CD8^+^ T cells, and NK-like T cells, can indeed belong to the same clone despite their diverse differentiation states (Figure S5D); yet cells from all three populations were consistently grouped within the same clonal niche in each clone (Figure S5E). Thus, clonal organization captures an axis of T-cell heterogeneity that is only partially aligned with conventional differentiation states, with cells of the same differentiation state distributed across multiple functional niches and individual niches composed of multiple different differentiation states.

Together, these analyses demonstrate that clonal organization is not simply a reflection of conventional T-cell differentiation states. While identity niches correspond to well-defined transcriptional states of T cells, functional niches reveal distinct clonal programs that go beyond differentiation states. Specifically, CloneTrast uncovers functionally distinct states of T-cell clones, demonstrating differences in cytotoxicity, tissue residency, stress responses, and other functional programs that cannot simply be recovered using standard expression-based representations.

### Applying CloneTrast to external test dataset of 1 million T cells

In order to test the generalization of our approach and its ability to capture a meaningful learned representation of T-cell clones across different datasets, we applied our pre-trained optimized model to additional test datasets composed of 11 different studies with paired scRNA/TCR-seq, consisting of 488 samples with more than 1 million T cells (Figure 4A, Table S1). These datasets, which also include two cancer types not represented in the training data, were entirely external to the model and were not used as part of the training and hyperparameter tuning procedures. Applying CloneTrast to this test dataset using scRNA-seq alone, resulted in a low-dimensional representation of T cells, with separation to major clonal niches and their relative positioning, similar to that observed using our training dataset (Figures 2A, 4B, S6A). The generated representation successfully brought together T cells based on their clonal identity (Figure 4C), with significant improvement of the patient clonal purity score compared to that achieved by expression-based representation (p-value = 2.47·10^-55^, Figure 4D, Table S3, Method details). Furthermore, the complementary model for prediction of T-cell clone size demonstrated a significant correlation between the predicted and the observed clone size of each T-cell clone in the test dataset (Spearman-r = 0.459, p < 2.23·10^-308^, Figure 4E, Method details).

**Figure 4.**
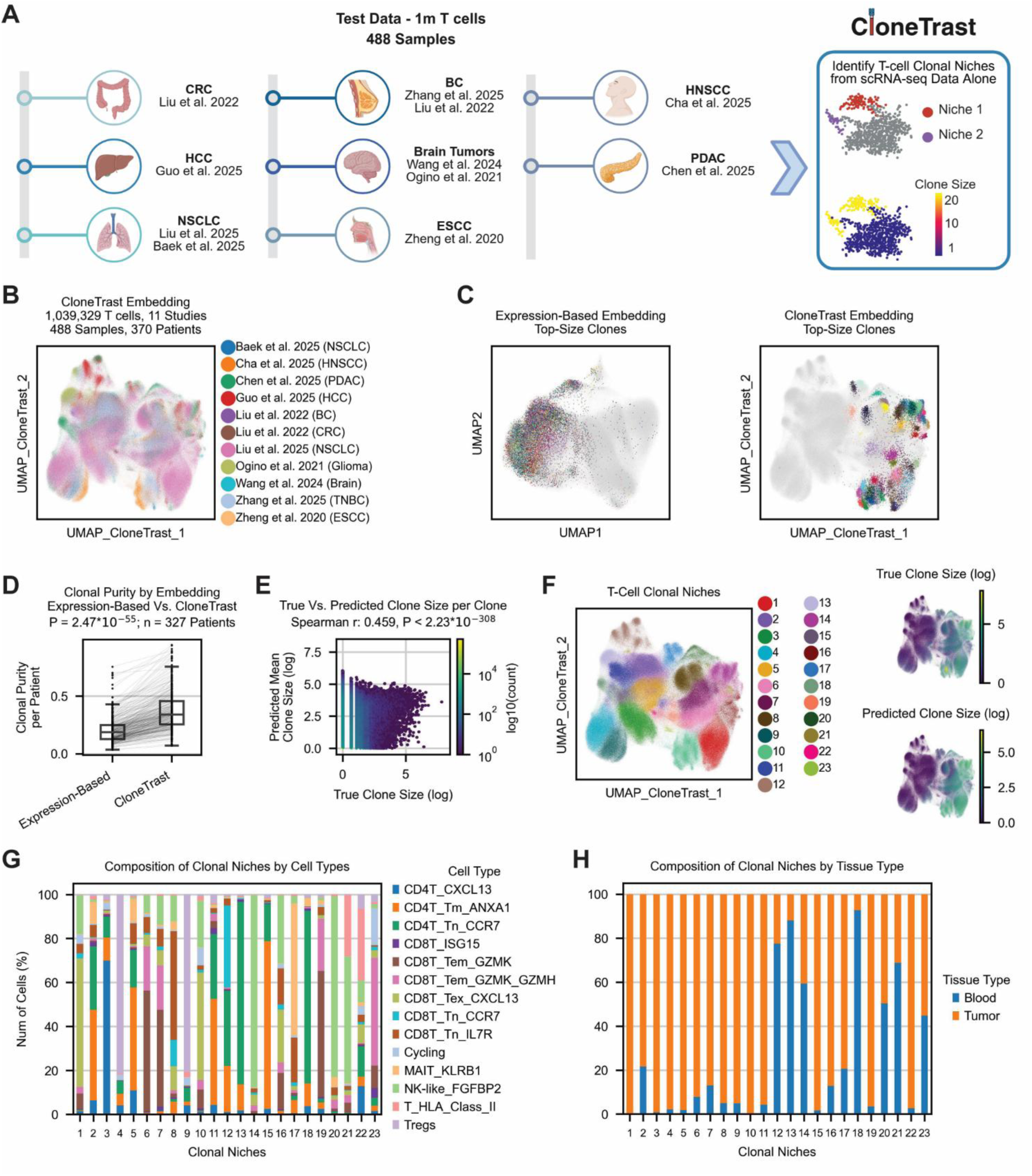
Applying CloneTrast to external test dataset of 1 million T cells. A. A schematic workflow depicting the application of the pre-trained model to external test datasets consist of more than 1 million T cells across 488 samples, spanning 11 different studies. B. Low-dimensional representation of the entire test dataset, obtained using CloneTrast, colored by 11 different studies. C. Expression-based representation (left) and CloneTrast-based representation (right), presenting T cells from top-size clones having clone size larger than 300 cells. The exact same clones are shown in both representations. D. Paired boxplots demonstrating the difference of clonal purity score per patient, calculated using either an expression-based embedding or the embedding obtained using CloneTrast (p-value = 2.47·10^-55^, n = 327, paired two-sided Wilcoxon signed-rank test). Only patients having clones with a minimal size of 20 cells were used for this calculation. E. Scatter plot showing the Spearman correlation between the observed clone sizes and those predicted by our model (Spearman-r = 0.459, p-value < 2.23·10^-308^). F. Leiden clustering results representing 23 distinct T-cell clonal niches upon the learned representation of CloneTrast (left), including overlay of both the observed and the predicted clone size for each T cell (right). G. Stacked barplot showing the composition of each clonal niche by different T-cell types. H. Stacked barplot showing the composition of each clonal niche by tissue type of origin, including tumor and blood samples. See also Figure S6 as well as Tables S1 and S3.

We next clustered T cells within the learned representation using the Leiden algorithm^68^ (Figure 4F, Method details). This step resulted in 23 distinct clusters of T cells reflecting different clonal niches with varying levels of clone sizes, as well as different composition of T-cell states, spanning both tumor and blood samples (Figures 4F-H and S6B-C, Table S3). While the identified clonal niches observed for the test dataset are not fully identical to those observed in the training dataset, the cellular composition of certain niches demonstrates major similarities between both datasets, as reflected by their total variation distance (TVD, Table S3, Method details). Reviewing niche composition, we found that multiple differentiation states span several clonal niches, recapitulating the patterns observed in the training data. This includes CXCL13-expressing CD8^+^ T cells, NK-like T cells, GZMK- and GZMH-expressing effector-memory CD8^+^ T cells, and cycling T cells (Figure 4F-H). In contrast, other differentiation states such as Tregs, CXCL13-expressing CD4^+^ T cells, and MAIT cells, are all located within distinct clonal niches, often within one or two niches (Figure 4F-H). Together, these findings demonstrate that the pre-trained CloneTrast model generalizes to independent datasets, including cancer types not represented during training, while recapitulating key relationships between T-cell differentiation states and clonal niches, as similarly observed in the training data.

### Pan-cancer clonal niches of T cells are associated with patient response to therapy

To examine the biological significance of the clonal niches captured by CloneTrast, we tested the association of different clonal niches with patient clinical outcomes and examined their dynamics following treatment, as well as between tumor and blood samples. We focused on 466 tumor samples from the training data, obtained from 237 patients and representing 17 different studies of cancer patients treated with immunotherapy across the training data^12,13,31,35–40,42–44,46–48,51^ (Figure 5A, Table S1).

**Figure 5.**
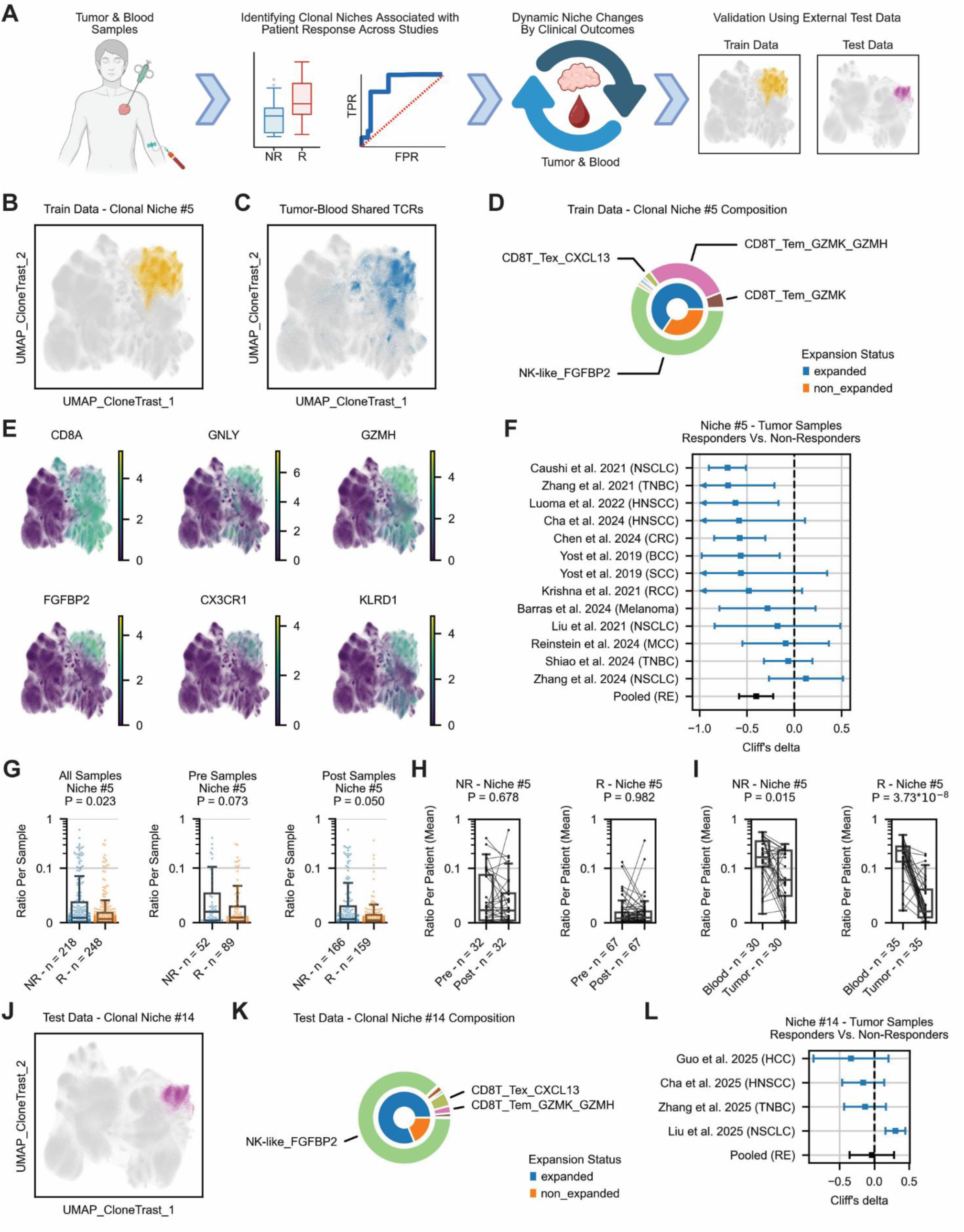
Pan-cancer clonal niches of T cells are associated with patient response to therapy. A. A schematic workflow depicting the quantification of clonal niche abundance across tumor and blood samples of cancer patients, demonstrating their association with patient clinical outcomes and dynamics following treatment. B-C. Low-dimensional representation of the training dataset based on CloneTrast, labeled with T cells from niche #5 (B) and T cells having TCRs shared between tumor and blood samples (C). D. Double donut chart representing the cell type composition of niche #5 (outer ring), as well as the composition of T cells according to their clonal expansion status (inner ring). E. Low-dimensional representation of the train dataset based on CloneTrast, demonstrating gene expression of selected markers, including *CD8A*, *GNLY*, *GZMH*, *FGFBP2*, *CX3CR1*, and *KLRD1*. F. Forest plot showing study-level Cliff’s delta effect sizes^76^ (blue), as well as pooled effect estimate (black), for the association between niche #5 abundance and patient response across independent studies. Squares represent study-level effect size, with horizontal lines indicating 95% confidence intervals. The pooled random effects were obtained using the DerSimonian-Laird method^74^. The vertical dashed line indicates no difference between both response groups. Positive values indicate higher abundance in responders, whereas negative values indicate higher abundance in non-responders. Only studies containing at least two responders and two non-responders were included, resulting in eligible 13 out of the 17 studies with available response annotations. G. The fraction of cells from clonal niche #5 per sample, compared between responding and non-responding patients across all tumor samples from all the 17 studies (left, n = 466), as well as a separate comparison by treatment time-points, including either pre- or post-treatment samples (right, n = 141 and 325, respectively). P-values were calculated using a two-sided Wilcoxon rank-sum test. H. Changes of niche #5 abundance following therapy in non-responders (left, n = 32), and responders (right, n = 67). P-values were obtained using a paired two-sided Wilcoxon signed-rank test. I. Changes of niche #5 abundance between tumor and blood samples in non-responders (left, n = 30), and responders (right, n = 35). P-values were obtained using a paired two-sided Wilcoxon signed-rank test. J. Low-dimensional representation of the test dataset based on CloneTrast, including labeling of T cells from niche #14. K. Double donut chart representing the cell type composition of niche #14 (outer ring), as well as the composition of T cells according to their clonal expansion status (inner ring). L. Forest plot showing study-level Cliff’s delta effect sizes (blue), as well as pooled effect estimate (black), for the association between niche #14 abundance and patient response across the four independent studies from the test dataset. See also Figure S7 as well as Tables S1 and S4.

We quantified the abundance of every clonal niche in each tumor sample, and compared these abundances between responders and non-responders (Method details). A random-effects meta-analysis using the DerSimonian-Laird method^74^ revealed two clonal niches, #4 and #5, that are significantly associated with patient response, such that both are consistently more abundant in non-responders across studies, with niche #5 showing the strongest signal (Table S4, Method details). While niche #4 is composed primarily of non-expanded CD4^+^ T cells across multiple differentiation states (Figures 2G, S7A-C, Table S4), niche #5 contains a substantial fraction of expanded T cells, with a majority of NK-like T cells expressing *FGFBP2*, *GNLY*, *GZMH*, and *CX3CR1*. Niche #5 also includes various fractions of CD8T_Tem_GZMK_GZMH, CD8T_Tem_GZMK, and CD8T_Tex_CXCL13 cells. Interestingly, this niche was found to be the most enriched with T cells sharing TCRs between tumor and blood samples among all clonal niches (32.61% of the cells in that niche, Figure 5B-E, S7A-B). Moreover, the association of niche #5 with clinical outcomes was consistent across studies despite substantial data heterogeneity in cancer types, treatment regimens, sampling time, and sample sizes (Figure 5F, Table S4, Method details). This signal was found in individual datasets, and was also observed when pooling all tumor samples together (p-value = 0.023, n = 466), and when separating them into pre- (p-value = 0.073, n = 141) and post-treatment samples (p-value = 0.05, n = 325, Figure 5G).

When considering only paired longitudinal biopsies with both pre- and post-treatment tumor samples, we found no change in niche #5 abundance between time points (Figure 5H, Method details). However, comparing paired tumor and blood samples, we found significantly lower abundance of this niche in tumors compared to their matched blood samples, both in responders and non-responders (p = 3.73·10^-8^ and 0.015 for responders and non-responders, respectively, Figure 5I, Method details). This finding suggests that this niche is largely represented in circulation, while its tumor-associated abundance is linked to poor clinical outcomes and remains stable across treatment time points. A similar observation was previously reported in melanoma patients, showing that blood-borne clonotypes from adoptive cell therapy products (ACTPs) repopulated mostly blood in responders, while infiltrating tumors of non-responders^75^. Notably, although niche #5 contains high number of NK-like T cells, when performing this analysis at the level of individual T-cell differentiation states, regardless of clonal niche identity, NK-like T cells showed no significant difference in their abundance between responders and non-responders (Figure S7D, Table S4, Method details). This result suggests that the association with patient response is niche-specific and cannot be explained solely by the abundance of the individual differentiation states composing each niche.

A complementary pattern was observed for CXCL13-expressing CD8^+^ T cells. Whereas the abundance of clonal niches enriched for these cells was not significantly associated with patient response (Table S4), CXCL13-expressing CD8^+^ T cells considered as a whole, irrespective of clonal niche identity, was significantly more abundant in responders (Figure S7E, Table S4), consistent with previous studies^14,69^. Thus, while the association of NK-like T cells with patient response was revealed only within a specific clonal context, the association of CXCL13-expressing CD8^+^ T cells was evident at the broader differentiation-state level. These findings suggest that clonal niche identity and conventional differentiation states capture complementary aspects of T-cell biology and its association with immunotherapy response.

We next focused on our test datasets, containing 345 tumor samples from 297 patients, representing 4 different studies having response annotations across the test data (Tables S1 and S4, Method details). Examining the corresponding NK-like T-cell niche in this data, niche #14, we found that in similar to the observation in the training data, this niche is enriched with T cells sharing TCRs between tumor and blood samples (49.56% of the cells in that niche, Figure 5J-K). Moreover, a similar trend towards abundance in non-responders was observed in 3 out of the 4 datasets, with the largest effect observed in hepatocellular carcinoma (HCC) patients^30^ (Figure 5L, Table S4). Taken together, these results show that clonal niche identity can reveal associations with therapy response that are not captured by conventional T-cell differentiation states alone, with related clonal patterns also observed across independent test datasets.

### Blood-borne and tumor-resident antigen-specific T cells are preferentially associated with distinct clonal niches

Given the high TCR diversity within each clonal niche (Figure S2B-C), we asked whether these niches exhibit selective enrichment for functionally distinct subsets of T-cell clones based on antigen specificity. To address this question, we first analyzed a dataset from our test cohort having paired scRNA/TCR-seq of HBV^+^ HCC patients receiving neoadjuvant anti-PD1 plus Lenvatinib^30^. This dataset includes HBV-dextramer-barcoded single-cell sequencing information across tumor and blood samples of five different patients, enabling the identification of HBV-specific T cells based on actual antigen binding. This dextramer panel comprised 31 experimentally validated HBV peptides known to induce CD8^+^ T cell responses, covering the core, envelope, polymerase, and X proteins^30^, used for sorting of dextramer^+^ CD8^+^ T cells for downstream sequencing. We identified and labeled 19 high-avidity HBV-specific T-cell clones across tumor and blood samples, following a pre-processing approach similar to that originally described^30^ (Figure 6A, Method details). Of note, although several of these 19 high-avidity clones were found to be specific to the same HBV antigen, they all had different TCR sequence identities. When examining their distribution across clonal niches, we found that these clones were preferentially mapped to niche #14, which contains the largest fraction of tumor-blood shared TCRs among all other clonal niches (Figure 6B-F), and a high number of expanded NK-like T cells (Figures 5J-K, 6F-G). Thus, the selective enrichment of these HBV-specific T-cell clones within this niche, whose specificity was established independently of the model, supports the biological relevance of the clonal-functional organization captured by CloneTrast. Importantly, the entire NK-like T cell population was distributed across multiple other niches (Figures 4G, 6G), highlighting the ability of CloneTrast to go beyond expression-based T-cell populations, and capture specific clonal niches with distinct biological properties.

**Figure 6.**
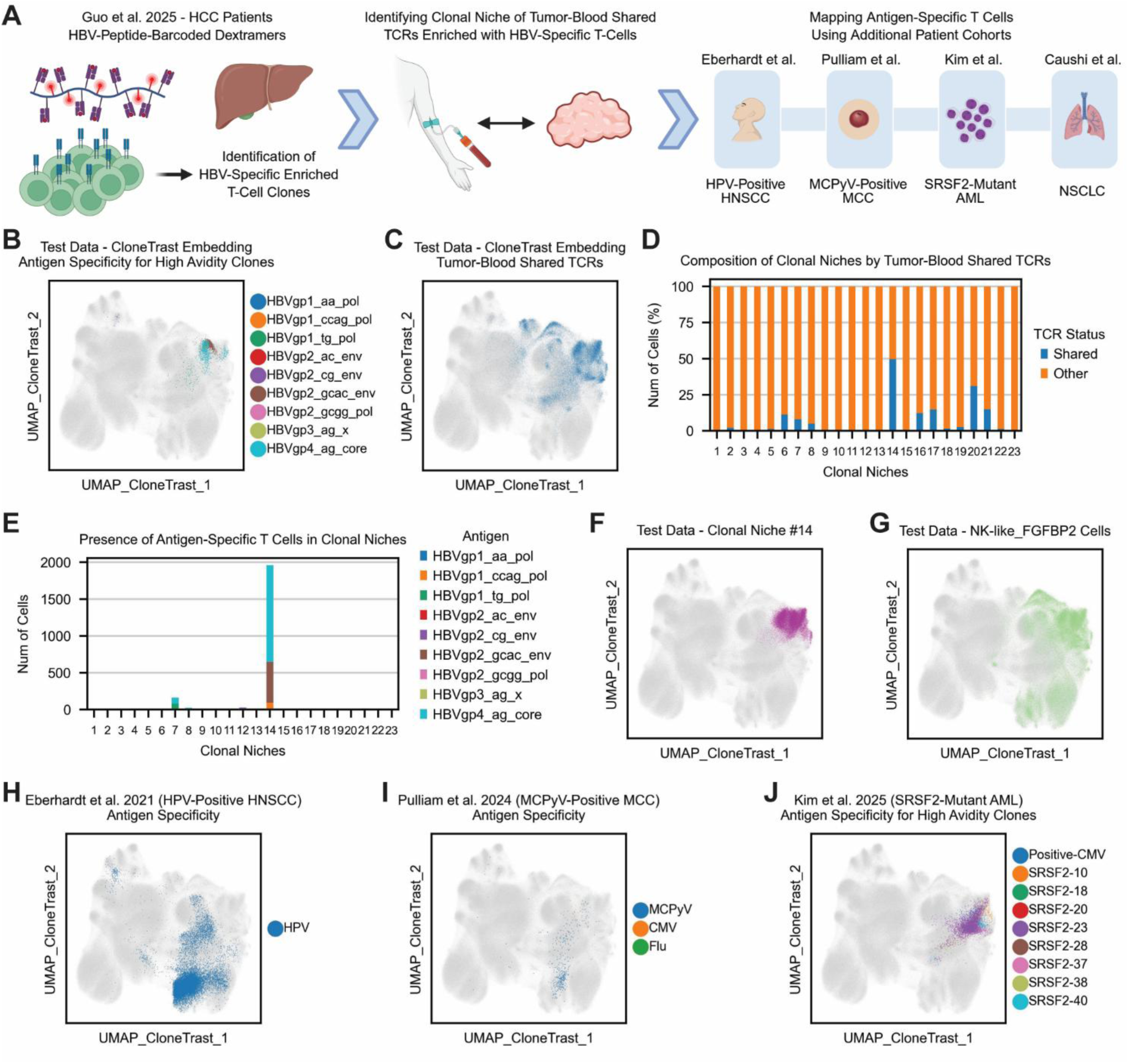
Blood-borne and tumor-resident antigen-specific T cells are preferentially associated with distinct clonal niches. A. A schematic workflow depicting the identification of high-avidity HBV-specific T-cell clones across five HBV^+^ HCC patients^30^, followed by detection of their presence in a clonal niche of tumor-blood shared TCRs. Additional cohorts with labeling of antigen-specific T cells were used to map the antigen-specific clonal landscape. B-C. Low-dimensional representation of the entire test dataset, obtained using CloneTrast, including labeling of T cells based on their clonal antigen specificity (B), and T cells having tumor-blood shared TCRs (C). HBV-specificity is shown only for T cells from high-avidity clones. D-E. Composition of clonal niches across the test dataset, based on TCR-sharing between tumor and blood samples (D), as well as on clonal antigen specificity (E). HBV-specificity is shown only for T cells from high-avidity clones. F-G. Low-dimensional representation of the test dataset based on CloneTrast, including labeling of T cells from niche #14 (F), and NK-like T cells (G). H-J. Low-dimensional representation of the test dataset based on CloneTrast, including labeling of antigen-specific T cells from the projected datasets of Eberhardt et al.^32^ (H), Pulliam et al.^33^ (I) and Kim et al.^34^ (J). See also Figure S8 and Table S1.

In order to further explore the clonal-functional landscape of antigen-specific T cells, we analyzed two additional scRNA/TCR-seq datasets of virus-driven malignancies (Table S1, Method details), including tumors of patients with HPV-positive head and neck squamous cell carcinoma (HNSCC)^32^, as well as tumor and blood samples of patients with advanced Merkel cell polyomavirus (MCPyV)-positive Merkel cell carcinoma (MCC)^33^. Both datasets used barcoded MHC-I multimers to identify HPV- and MCPyV-specific T cells, respectively. We applied CloneTrast to these datasets and projected their antigen-specific T cells upon the low-dimensional representation obtained for the test data (Figure 6H-I, Method details). In both cases, antigen-specific tumor-infiltrating T cells were predominantly mapped to clonal niche #10, a niche enriched for CXCL13^+^ CD8^+^ T cells in which 99.46% of the TCRs were not shared between tumor and blood samples (Figure 6D, Figure S8A-C). Thus, HPV- and MCPyV-specific T cells converged on a shared clonal-functional niche with limited tumor-blood TCR sharing, distinct from the niche preferentially occupied by HBV-specific clones. To go beyond virus-driven malignancies, we analyzed an additional scRNA/TCR-seq dataset of blood samples from patients with acute myeloid leukemia (AML) harboring mutations in the splicing factor *SRSF2*^34^ (Table S1). We used the dextramer panel of 46 mis-splicing-derived neoantigens provided by the authors to label neoantigen-specific high-avidity T-cell clones (Figure 6J, Method details). High-avidity T-cell clones from these blood samples were predominantly mapped to two adjacent clonal niches, including niches #6 and #14 (Figure S8D). Similarly, mutation-associated neoantigen (MANA)-specific T cells from non-small cell lung cancer (NSCLC) tumors^31^ were predominantly mapped to a clonal niche enriched with CXCL13^+^ CD8^+^ T cells (Figure S8E-F, Method details). Together, these findings indicate that antigen-specific T cells occupy distinct clonal-functional states with their tissue context. Tumor-infiltrating HPV-, MCPyV-, and MANA-specific T cells were preferentially mapped to a CXCL13^+^ CD8^+^ T-cell-enriched niche characterized by limited tumor-blood TCR sharing, whereas blood-derived AML neoantigen-specific clones were mapped predominantly to niches #6 and #14, the latter also enriched for HBV-specific clones and tumor-blood shared TCRs. These observations support the ability of CloneTrast to capture biologically meaningful organization among antigen-specific T cells across cancer types and antigen classes. Establishing the clinical significance of these clonal-functional states in the context of antigen specificity will require larger cohorts integrating clinical outcomes, antigen-binding assays and paired scRNA/TCR-seq from matched tumor and blood samples.

## Discussion

The functional organization of T-cell clones is central to understanding anti-tumor immunity, yet it remains challenging to study across large single-cell datasets because clonal information is typically derived from paired TCR sequencing. Here, we present CloneTrast, a supervised contrastive-learning framework that uses paired scRNA/TCR-seq data during training to learn a transcriptome-based representation of T-cell clonal function. In contrast to existing methods designed primarily for joint integration of paired gene-expression and TCR sequence modalities, CloneTrast uses clonal identity as supervision during training but does not require TCR sequence information at inference. This enables the application of a pre-trained model to scRNA-seq datasets in which paired TCR sequencing is unavailable, incomplete, or sparsely captured.

Using a large human pan-cancer database of paired scRNA/TCR-seq profiles from tumor and blood samples, CloneTrast identified groups of T-cell clones with shared transcriptional-functional properties, which we term ‘clonal niches’. These niches represent an intermediate level of T-cell organization: they are distinct from individual clonotypes defined by identical TCR sequences, but also differ from conventional T-cell state annotations derived from standard transcriptomic clustering. While some clonal niches corresponded closely to canonical T-cell states, such as Tregs, naive T cells, and MAIT cells, others were composed of multiple annotated T-cell states. Thus, CloneTrast captures functional relationships between T-cell clones that are not fully resolved by standard single-cell workflows.

A key feature of CloneTrast is that it learns from clonal identity without using the actual TCR sequence as a model input. This distinction is important because it allows the model to identify transcriptionally similar clonal programs even among T-cell clones with different receptor sequences. Consistent with this, GLIPH2-based analysis showed that clonal niches are not simply explained by CDR3 sequence similarity. Although grouping TCRs by sequence similarity reduced the measured diversity within niches, diversity remained high overall, supporting the notion that CloneTrast captures convergent transcriptional states of T-cell clones rather than merely grouping cells by related TCR sequences. Thus, the low-dimensional representation of CloneTrast has two complementary properties: (1) it captures the clonal structure of T cells within patients, as demonstrated by the high clonal purity scores, and (2) it identifies broader clonal niches that reflect clonal-functional organization across patients and studies, at the pan-cancer level.

Beyond identifying clonal niches, CloneTrast provides an interpretable framework for characterizing the transcriptional features underlying niche identity. Using model explainability, we identified genes that contribute to the positioning of cells within specific clonal niches across the learned representation. This approach goes beyond standard differential expression analysis, because genes that distinguish a niche transcriptionally are not necessarily the same genes that drive its organization in the latent space. In niches dominated by canonical T-cell states, explainable features often reflected expected marker genes. In more heterogeneous niches, however, explainable genes highlighted more complex programs that may underlie clonal-functional differences between transcriptionally similar populations. This was illustrated by CXCL13⁺ CD8⁺ T cells, which were separated by CloneTrast into multiple clonal niches with distinct transcriptional features, despite appearing more overlapping upon an expression-based representation.

A specific niche enriched with TCRs shared between tumor and blood samples, having high numbers of NK-like T cells, was associated with patient response to immunotherapy across studies, being significantly more abundant in tumors of non-responding patients. Notably, this association was not observed by NK-like T cells as a population, but rather by a subset of these cells that existed in that specific niche. This supports the idea that clonal niches capture functional information that is not fully represented by conventional cell-state labels alone. Importantly, this association with clinical outcomes was observed despite the heterogeneity of cancer types, treatment regimens, response definitions, sampling time points, and cohort sizes across the integrated datasets. It is important to note though that certain individual studies showed contradicting trends^46,62^.

The analysis of HBV-positive hepatocellular carcinoma cohort further demonstrated how CloneTrast can recover biologically meaningful antigen-related clonal structure. Although CloneTrast does not use TCR sequence or antigen specificity as input during inference, high-avidity HBV-specific T-cell clones were enriched within a defined clonal niche that also contained a high abundance of TCRs shared between tumor and blood samples. This finding suggests that clonal niches can capture functionally coherent T-cell populations related to antigen-driven biology. Furthermore, analysis of additional patient cohorts with antigen-specific T cells, spanning different cancer types and antigens, revealed that tumor-resident and blood-borne antigen-specific T cells are preferentially associated with distinct clonal niches. Together, these findings support the ability of CloneTrast to recover biologically meaningful clonal organization linked to antigen specificity. However, one should not interpret clonal niches as being exclusively composed of tumor-specific T cells, as non-tumor-specific T cells may also be present in the same niches, sharing common transcriptional and functional characteristics with tumor-specific T cells. Indeed, the potential contribution of non-tumor-specific intra-tumoral T cells to tumor-killing is increasingly being studied^77,78^.

Several limitations should be considered. First, CloneTrast was trained and evaluated using tumor and blood samples from human cancer patients, and its generalizability to other biological contexts, including normal tissues, lymphoid tissues, autoimmune diseases, infection, or non-human systems, remains to be tested. Second, the identified clonal niches depend on the learned representation and on the clustering strategy applied to it; different clustering resolutions may reveal broader or finer levels of clonal-functional organization. Third, although CloneTrast can infer clonal-functional states from scRNA-seq alone, it does not infer TCR sequence, clonotype identity, or antigen specificity directly. Thus, conclusions regarding antigen specificity require independent validation using paired TCR sequencing, antigen-binding assays, or other orthogonal approaches. Finally, clinical associations identified across heterogeneous public datasets should be considered hypothesis-generating and will require validation in larger, more uniformly treated patient cohorts.

Overall, CloneTrast provides a scalable and interpretable framework for studying the clonal-functional landscape of T cells from single-cell transcriptomes. By learning from paired scRNA/TCR-seq data, but enabling inference using scRNA-seq alone, CloneTrast extends clonal-level analysis to datasets in which direct TCR information is unavailable or incomplete. This approach reveals pan-cancer T-cell clonal niches associated with functional states, antigen-related biology, and patient response to therapy, offering a new computational strategy for dissecting T-cell clonal organization in cancer.

## Resource availability Lead contact

Further information and requests for resources should be directed to and will be fulfilled by the lead contact, Keren Yizhak.

## Materials availability

This study did not generate new unique reagents.

## Data and code availability

- This paper analyzes existing, publicly available scRNA/TCR-seq data. The accession numbers for these datasets are summarized in Table S1.
- All original code has been deposited at https://github.com/yizhak-lab-ccg/CloneTrast and is publicly available as of the date of publication. The complete set of trained models is publicly accessible via Figshare^79^ (https://doi.org/10.6084/m9.figshare.32228991). The full documentation of our platform, including tutorials and guidelines, is available on Read the Docs (https://clonetrast.readthedocs.io). An official Python package is provided via PyPi (https://pypi.org/project/clonetrast).
- Any additional information required to reanalyze the data reported in this paper is available from the lead contact upon request.

## Acknowledgments

This work was supported by the Israel Science Foundation (1124/25), the Israel Cancer Research Fund (23-204-RCDA) and the Career Advancement Chairs in Artificial Intelligence – Schmidt Futures. This work received additional support by the Ruth and Bruce Rappaport Technion Integrated Cancer Center (RTICC). Figures 1A-C, 3A, 4A, 5A, and 6A, were created with BioRender.com using a paid license.

## Author Contributions

O.S., A.P. and K.Y. conceived the idea. O.S. and K.Y. designed the study. O.S. created the code repository.

O.S. performed the analysis. O.S. and K.Y. wrote the manuscript. O.S., A.P. and R.A. reviewed the code repository. O.S., A.P., R.A. and K.Y. reviewed the final version of the manuscript.

## Experimental model and study participant details

This study did not involve any new experimental models, human participants, or animal subjects. All data used in this study are publicly available, with samples and single cells obtained from each dataset described in Table S1 following our quality control. Details regarding data acquisition and processing are provided in the method details.

## Method details

### Acquisition of paired scRNA/TCR-seq datasets and preprocessing

26 single-cell datasets of cancer patients having paired scRNA/TCR-seq data were collected together with their annotations for clinical outcomes and treatment time-point, where available. 15 datasets contained tumor biopsies, 9 datasets contained both tumor and blood samples, and 2 datasets contained blood samples alone (Table S1).

All scRNA-seq datasets were droplet-based and contained unique molecular identifier (UMI) counts. For each dataset, we followed similar preprocessing steps to those we previously conducted^15,21^ using Scanpy^80^: we removed cells expressing less than 200 genes and removed genes that were expressed in less than 3 cells. We also removed cells having more than 10% of mitochondrial gene-count. In the dataset of Krishna et al.^36^, scRNA-seq was provided following the authors’ original quality-control (QC) while excluding mitochondrial genes from the provided gene expression matrix. In that case only, we used the expression matrix as provided by the authors following their QC, retaining cells with less than 20% of mitochondrial gene-count. We also applied Scrublet^81^ to remove potential doublets and filtered-out cells having *‘doublet_score’* larger than 0.3 and/or those predicted as doublets with score below 0.3.

For the scTCR-seq datasets, we rearranged each dataset to be compatible with the Adaptive Immune Receptor Repertoire (AIRR) schema^82,83^ that was further preprocessed with Scirpy^84^ and could be directly uploaded using the *‘scirpy.io.read_10x_vdj’* function. For each scTCR-seq dataset, we applied quality control based on the standard protocol suggested by the authors. In short, we removed cells having multi-chains, orphan VJ or orphan VDJ chains. Following this quality control process, we removed samples having less than 3 cells overall, as well as 18 single cells having extreme CDR3 sequence lengths above 30 amino acids, and integrated all processed datasets resulting in a total of 1,055,915 single cells from tumor samples and 532,376 single cells from blood samples, all with paired scRNA/TCR-seq. Overall, 11,950 genes existed across all datasets and were used for further analysis. Importantly, as we used the original clinical metadata provided by the authors for each dataset, patient response was defined differently across studies, whenever existed, and was provided mostly by using either RECIST^85^ or pathological response annotations. In addition to the differences between both response definitions, in certain studies using pathological response alone, definitions were inconsistent either, making patient clinical outcomes highly heterogeneous. For consistency and simplicity purposes, we labeled samples having RECIST complete/partial response (CR/PR) as responders and those with stable/progressive disease (SD/PD) as non-responders, as we similarly did previously^8,15^. We also considered responders as those with high/major/complete pathological response and non-responders with non-high/major/complete pathological response, as was done previously^15^. In the dataset of Luoma et al.^37^, response was provided both by RECIST and as pathological response. In that case, and because many patients had a ‘not measurable’ response according to RECIST but did have pathological response, we considered responders as those with high pathological response and non-responders with a non-high pathological response.

Overall, although not fully aligned, these coarse response definitions enabled us to robustly evaluate general associations with clinical outcomes following model application, as described below. Importantly, these associations should be refined as more uniform clinical data become available.

### Markov Affinity-based Graph Imputation of Cells (MAGIC) for detection of drop-outs and clone labeling

In order to identify possible sequencing drop-outs in our integrated dataset, we first normalized the expression level to a standard target sum of 10,000 counts per cell and then applied log-transformation. We applied MAGIC^86^ to the log-normalized count matrices of tumor and blood samples separately, using all the genes and single cells passing our QC, while using the default arguments of the algorithm implementation in Scanpy^80^. We first fitted a density curve to the log-transformed imputed gene expression of *CD8A/B*, *CD4*, and *FOXP3*, and set the expression threshold for each gene as the trough of the bimodal density curve. For the imputed gene expression of *FOXP3*, the threshold was set as 1. For the non-imputed expression of these genes, expression threshold was also set as 1. Overall, all the T cells that passed our QC process were further divided into five subtypes (Figure S1A-E): we considered a single cell to be CD8^+^, when the imputed or non-imputed gene expression of either *CD8A* or *CD8B* was above the threshold. Similarly, a single cell was considered to be CD4^+^, when the imputed or non-imputed gene expression of *CD4* was above the threshold. We also defined double-positive cells as those that were considered to be both CD8^+^ and CD4^+^. Double-negative cells were those that were both CD8^-^ and CD4^-^. Tregs were defined as those that are CD4^+^ having imputed or non-imputed gene expression of *FOXP3* above the threshold. In cases where the imputed expression resulted with loss of signal from the non-imputed gene expression, we remained consistent with the non-imputed gene expression. Importantly, the imputed gene expression was used solely for cell type classification, while the model input itself included non-imputed gene expression to retain the original data structure and prevent potential artifacts introduced by imputation.

We then defined clonotypes based on the identity of the CDR3 nucleic acid sequence of each cell using *‘scirpy.tl.define_clonotypes’*, such that both the VJ and VDJ CDR3 sequences had to match and the T-cell subtype of all the cells in each clone is the same (i.e. CD8^+^, CD4^+^, double-positive, double-negative or Treg). In cases where more than one pair of VJ and VDJ sequences was detected per cell, we considered the most abundant pair of VJ/VDJ chains where applicable. In the dataset of Krishna et al.^36^, scTCR-seq was provided only with the amino acid sequence of the CDR3 region. In that case only, clonotypes were defined based on the amino acid identity of the CDR3 sequence. Of note, an in-house evaluation using a more conservative clonotype definition, requiring both identical CDR3 sequences and identical V genes (using the argument ‘*same_v_gene = True*’), resulted with 99.78% concordance of clonotype identity compared to using CDR3 sequences alone. Similar concordance was achieved by requiring identical J genes (using the argument ‘*same_j_gene = True*’), in addition to identical CDR3 sequences and identical V genes. To ensure our framework is robust and compatible with studies reporting only CDR3 sequences, we based clonotype definitions solely on CDR3 identity. This process resulted in categorical clone identities (e.g. clone A, clone B, clone C, etc.), that were later used for supervision during model training, rather than feeding actual TCR sequence identities to the model. Lastly, in order to obtain a simple binary definition of clonal expansion, we defined a clonotype as ‘expanded’ per sample according to Shiao et al.^39^, and as we did previously^15,21^, such that ‘expanded clones’ were those that contained more than 1.5x the median number of cells found in each clonotype in that sample, while excluding singletons. This approach normalizes for differences in sampling depth and repertoire complexity between samples and avoids the biases introduced by fixed-clone-size thresholds, enabling consistent identification of clonal expansion across datasets. This relatively conservative threshold reduces the impact of noise while capturing clones with potential functional impact. Importantly, this definition was used solely for niche-characterization purposes (Figures 5D, 5K, S6C, S7A), with no actual influence on the downstream analysis.

### Integration of scRNA-seq data and cell type annotations

In order to integrate our scRNA-seq data, we normalized the expression level in the integrated scRNA-seq dataset to a standard target sum of 10,000 counts per cell and then applied log-transformation. Highly variable genes were calculated using *‘scanpy.pp.highly_variable_genes’* with a *‘batch_key’* of *‘study’* in order to avoid selection of study-specific genes, ensuring better downstream batch correction. We then calculated a 50-component PCA and applied Harmoy^87^ with a ‘*batch_key’* of *‘sample’* in order to remove batch effects between samples to obtain Harmony-corrected PCA results. We used ‘*scanpy.pp.neighbors*’ over the Harmony-corrected PCA to obtain a neighbor graph, followed by UMAP^88^ for dimensionality reduction and data visualization. We applied the unsupervised Leiden^68^ clustering algorithm with a resolution of 1.5 in order to obtain fine-grained T-cell clustering, and calculated differentially expressed genes for each cluster using the ‘*scanpy.tl.rank_genes_groups*’ function with the Wilcoxon rank-sum test (Table S2). Following, we manually annotated cell types according to the top differentially expressed genes of each cluster, and used the publicly available cell type markers that were provided by Liu et al.^62^ as an additional reference for T-cell subtype identity. Across all the 1.5 million T cells that were annotated, 646 cells had an expression of ambiguous gene markers and were consequently labeled as ‘unassigned’ cell type, however these cells were included as part of the model input during training using their clonal identities. Importantly, this entire pipeline was preformed solely for the purpose of annotating cell types and creating a standard expression-based representation of the cells, while the model itself was trained using all the 11,950 genes in the data (and not just the highly variable genes), as described below.

### CloneTrast model design

We implemented CloneTrast as a feed-forward expression encoder trained with supervised contrastive loss. For each cell, preprocessed row-normalized and log-transformed gene expression x ɛ R^n_genes^ is passed through an encoder, to obtain a latent embedding h. A projection head maps h to a contrastive space z, where training is performed using a SupCon objective, as described below. For each training batch, positive pairs were defined as cells sharing the same categorical clone identity (‘*clone_id*’) per patient (e.g.clone A, clone B, clone C, etc.), rather than actual TCR sequence identities, while negatives are all the other cells in the batch. At inference, only the encoder embedding h is retained as the final model representation. To improve robustness to inter-patient effects, training used patient-wise batching such that each batch is formed by sampling cells from up to k patients, while still optimizing clone-level contrastive separation. This encourages the model to learn clone-discriminative structure both within and across patients, rather than overfitting to patient-specific expression signatures and improving batch effect removal between patients during training. As a complementary task, we trained a separate clone-size regressor (independent encoder) to predict log-clone-size from expression using a per-cell MSE loss combined with clone-level pairwise logistic ranking loss. The architecture of both the supervised contrastive model and the clone-size regressor were defined using the PyTorch^73^ library and included fully connected layers incorporating dropout regularization to mitigate overfitting, with an optimization handled using the AdamW optimizer^89^. Optuna^67^ was used for hyperparameter tuning as described below. Importantly, at inference, expression matrices are automatically aligned to the training gene set such that genes present in the query dataset but absent from training are removed, and genes present in training but missing from the query are imputed as zeros (thus mimicking sequencing drop-outs), while shared genes are reordered to exactly match training input order. This guarantees a fixed input dimensionality and consistency across datasets. Notably, the model was trained with gene representation using Ensembl IDs^90^ for generalization purposes and simpler user interface.

### Model training and optimization

CloneTrast was trained using a training data consisting of more than 1.5 million T cells (as described above), following a structured workflow, beginning with data preprocessing and splitting into training (80%) and validation (20%) sets. During data splitting we restricted all of the cells obtained from a single patient to be used in either training or validation sets, but not both, ensuring that data from the same patient cannot appear in both sets. Preprocessing included count row-normalization to a target sum of 10,000 counts per cell, followed by log transformation. Hyperparameter tuning was performed using Optuna^67^, which explores parameter spaces across multiple trials, integrating automated logging, checkpointing, early stopping, and systematic model evaluation to ensure reproducibility. During the optimization process we trained the model for 100 trials, such that the optimal hyperparameter configuration was tuned in order to maximize the validation set weighted per-patient clonal purity score across all trials. More explicitly, in every epoch for each patient *p* we:

1. Obtained the model embedding for that patient’s cells based on the current model weights.
2. Use that embedding with a cosine distance to find the 10 nearest neighbors of cells having clone size of at least 20 cells to avoid unstable estimates for small clones.
3. Calculate for each cell what fraction of its neighbors have the same ‘*clone_id*’.
4. Average these cell-level fractions per eligible clone (clones having at least 20 cells) to avoid over weighting of major clones.
5. Average the clone-level values per patient to get a patient purity score.

Finally, the overall clonal purity score was the average of all patient scores weighted by patient cell count (number of cells), so larger patients contributed proportionally more to the final optimization metric. This weighting was performed to reduce the effect of the high purity scores observed for patients having relatively lower amount of cells.

For each training trial during the optimization process, hyperparameters were tuned to maximize the overall clonal purity score, including tuning of the latent embedding size, contrastive space size, width and depth of the network, as well as the supervised contrastive loss variant used during model training, selected out of the two variants introduced by Khosla et al.^66^ 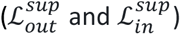.

More explicitly, for each anchor cell *i* in a batch, *P*(*i*) denotes the set of other cells in the batch that share the same clonal identity, and therefore serve as positive indices. *A*(*i*) denotes all other cells in the batch excluding the anchor itself, and therefore serve as negative indices. The projection vectors *z_i_* are L2-normalized, such that their dot product corresponds to cosine similarity. The temperature hyperparameter *τ* scales similarities:

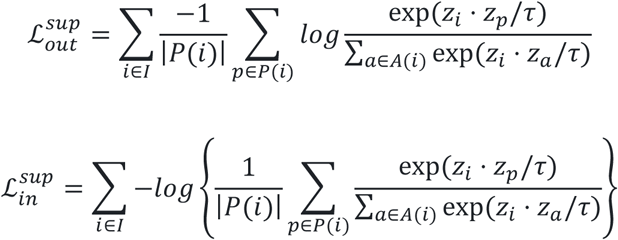

In 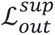 the logarithm is applied separately to each positive pair before averaging across positives, therefore encouraging every positive sample to have high similarity to the anchor. In 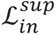 the positive similarities are averaged first, and the logarithm is then applied to that average. As a result, highly similar positives can compensate for other less similar positives, making 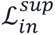 less sensitive to the transcriptional heterogeneity often seen among cells within the same clone. In fact, 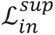 was the variant chosen for the optimized public pre-trained model by the automatic hyperparameter tuning process, resulting in higher clonal purity score.

For each training trial, the model was trained until no improvement of the overall clonal purity score was observed for 10 consecutive epochs. Overall, this entire process was conducted for 100 different trials, resulting in an optimal set of hyperparameters (Figure S2A). Importantly, the optimized trained model is compatible with Scanpy’s standard data structure^80^ and was made publicly available with support for automatic model download. Hardware used for optimization and training included a workstation equipped with an Intel core Ultra 9 285K CPU 256 GB RAM and an NVIDIA RTX 6000 Ada Generation GPU 48 GB VRAM. We applied a similar approach for the training and optimization of the clone size prediction model, such that the optimization objective was minimizing the sum of the cell-level and clone-level Wasserstein distances between the observed and the mean predicted log clone size of each clone. This sum also included one minus the Spearman r value between the predicted and the observed log clone sizes at the clone-level. For each training trial, the model was trained until no improvement of this objective was observed for 10 consecutive epochs.

The full lists of hyperparameters used during model training are described in Table S2, including both the contrastive and the clone size prediction models.

### Obtaining a low-dimensional representation and labeling of T-cell clonal niches

We applied the optimized trained model to the training data, obtaining a low-dimensional representation for each T cell. A k-nearest neighbor graph was then constructed in the embedding space using a cosine metric with 15 neighbors. This graph was then used both for visualization with UMAP, resulting in a two-dimensional representation of the data (Figure 2A), and for clustering using the Leiden algorithm with resolutions of 0.5, 0.75, and 1, to identify T-cell clusters, further termed as ‘clonal niches’. Following manual inspection of the clusters, the clustering resolution of 0.5 was chosen for downstream analysis (Figure 2E). In addition, we calculated the patient clonal purity score for the embedding obtained using CloneTrast (as described above), and compared it with an equivalent score obtained using a standard expression-based uncorrected 50-component PCA space, using a paired two-sided Wilcoxon signed-rank test (Figure 2C). Importantly, in order to make the patient clonal purity score comparable between CloneTrast and the PCA space, the 50-component PCA space in that case was calculated using the same genes that were used as an input for the model (instead of calculating PCA using just the most highly variable genes). Moreover, we evaluated the Spearman correlation between the observed log clone size and the mean predicted log clone size obtained using the complementary clone-size prediction model (Figure 2D).

### Application of GLIPH2 for clustering T-cell receptors

In order to examine whether clonal niches are associated with specific T-cell receptor identity patterns, we applied immGLIPH (https://github.com/BorchLab/immGLIPH), an R implementation of the GLIPH2 algorithm^71^ for clustering T-cell receptors. Rather than using identical CDR3 sequences for clonal identity, GLIPH2 identifies CDR3 specificity groups based on local (motif-based) and global (sequence-based) similarities, resulting in a more flexible grouping of T cells which is more permissive compared to strict sequence identity alone. We used our training dataset of 1.5 million T cells and prepared the relevant input for the immGLIPH implementation of the GLIPH2 algorithm using its default arguments, resulting in a new T-cell grouping solution (Figure S2B). In cases where a single T cell was attributed to multiple different specificity groups, it was ultimately classified into the largest of those groups. Following, we used the ‘*scirpy.tl.alpha_diversity*’ function with ‘*metric = normalized_shannon_entropy*’ in order to calculate the alpha diversity of GLIPH2 specificity groups per clonal niche, and used a paired two-sided Wilcoxon signed-rank test to compare these values with the alpha diversity of T-cell clones per clonal niche, obtained using identical CDR3 sequence identity, as described above (Figure S2C). It is important to note that immGLIPH currently supports only amino acid sequence of TCR beta chains, while we assigned clone identities using both CDR3α and CDR3β nucleic acid sequence identities. In addition, in order to include studies that do not report V gene annotations, we applied immGLIPH using solely CDR3 sequences.

### Application of mvTCR and MIST

In order to emphasize differences between the low-dimensional representation obtained using CloneTrast and that of existing scRNA/TCR-seq integration methods, we applied mvTCR^23^ and MIST^25^ using our training dataset of 1.5 million T cells. Importantly, while additional methods for joint analysis of paired scRNA/TCR-seq data do exist, we found mvTCR and MIST to be the only ones compatible with the memory efficient anndata^91^ structure, enabling relatively straightforward application to our training dataset, along with more scalable runtimes. More explicitly, we applied mvTCR according to the publicly available code provided by the authors, followed by extraction of the learned latent representation using the trained mvTCR model. Similarly, we applied MIST using the code provided by the authors and extracted its learned latent representation (Figure S2D). For both mvTCR and MIST, we used their built-in batch effect removal option, to remove batch effects between patients. Notably, while mvTCR requires the existence of both CDR3α and CDR3β amino acid sequences for each T cell, MIST additionally takes both the V and the J genes into account, thus limiting its applicability across datasets reporting CDR3 sequences alone.

Consequently, MIST’s automatic preprocessing excluded substantial amount of T cells from our training data prior to model training. Finally, we visualized the clonal niches identified by CloneTrast, across the learned representations obtained using mvTCR and MIST, further emphasizing the structural differences of each method (Figure S2E-G).

### Evaluation of batch effect removal across representations

We used the integration Local Inverse Simpson’s Index (iLISI) score, as implemented by the harmonypy package (https://github.com/slowkow/harmonypy)^87^, using the function ‘*compute_lisi*’. We applied this function to the two-dimensional UMAP representations obtained using multiple different embedding spaces. These included the embedding space obtained by mvTCR^23^, MIST^25^, CloneTrast, as well as that achieved by using a 50-component harmony-corrected, and an uncorrected PCA space (Figure S2H). As both PCA spaces were obtained using the top highly variable genes, we made an additional comparison to a UMAP representation of a 50-component PCA space obtained using all the available 11,950 genes in the training data (without filtering for highly variable genes). We considered this specific PCA space as a baseline of comparison. For each representation we measured the integration of samples by calculating the iLISI score of each T cell, and created a violin plot of these scores for each representation (Figure S2H, Table S2).

### Model explainability using Captum

To interpret niche-defining transcriptional features learned by CloneTrast, we performed an attribution analysis on single-cell gene expression profiles using Captum Integrated Gradients (IG)^72^. First, in order to define an attribution target, cells were projected to the model contrastive space, L2-normalized, averaged within each clonal niche using all the 18 clonal niches in the training data, and then L2-normalized again, to construct one L2-normalized prototype vector per niche. For each cell, we defined a differentiable niche-matching objective as the cosine similarity to its true niche prototype minus a smooth log-sum-exp term^92^ over similarities to non-matching niches (with temperature parameter *β* = 0.05), yielding a contrastive matching score.

Formally, we defined the niche matching objective as follows: Let cell *i* have the L2-normalized projection vector *z_i_* and belong to clonal niche *c*(*i*). Let *p_k_* denote the L2-normalized prototype of clonal niche *k*. The cosine similarity between cell *i* and niche prototype *k* is defined as: *s_i_*_,*k*_ = 〈*z_i_*, *p_k_*〉 = *z_i_* ⋅ *p_k_*. The contrastive match score for cell *i* is then defined as:

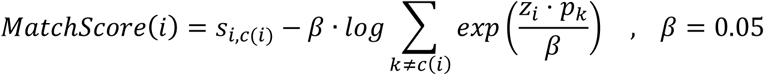

IG attributions were then computed with respect to this objective using 100 integration steps, with a baseline defined as the per-gene global mean expression across all single cells. Attributions were calculated per cell and then averaged by niche to obtain mean signed IG (directional contribution) and mean absolute IG (magnitude of contribution) for each gene. Genes were subsequently ranked within each clonal niche into top sets based on signed IG (Table S2). To improve directional interpretation, we also summarized the mean expression deviation from baseline and the sign of an approximate path-averaged gradient, that is, the gradient of the niche-matching score with respect to each gene, averaged over the integrated-gradients path, and then averaged across the cells within the niche. A positive sign of the path-averaged gradient indicates that higher expression of the gene increased the niche-matching score, whereas a negative sign indicates that lower expression increased the niche-matching score (Table S2). For visualization purposes only (Figure 3B), the top 15 ranked genes per clonal niche were used to demonstrate upregulation and downregulation patterns across clonal niches. Lastly, in order to demonstrate that CloneTrast captures clonal niche identities that go beyond simple gene expression patterns, we focused specifically on a population of CXCL13^+^ CD8^+^ T cells across five different clonal niches containing the largest count of cells from that population (Figure S3A-D). We performed differential gene expression analysis between these populations (using the ‘*scanpy.tl.rank_genes_groups*’ function, Figure S3D, Table S2), and applied a standard, expression-based dimensionality reduction process to the population of CXCL13^+^ CD8^+^ T cells alone, to show that its different niche identities cannot be recovered using a standard preprocessing pipeline (Figure S3E). We performed similar analysis for two additional populations of NK-like T cells (Figure S4A-C) and GZMK-expressing effector-memory CD8^+^ T cells (Figure S5A-C).

### Application of CloneTrast to external test data

In order to evaluate CloneTrast’s performance in an unbiased manner, as well as demonstrating its robustness and generalization, we applied the pre-trained optimized model to external test data containing 11 different studies having more than 1 million T cells across 488 tumor and blood samples (Figure 4A, Table S1). Importantly, these datasets were entirely external to the model and were not used during both training and optimization procedures. For each one of the different 11 studies, we applied a separate quality control using both data modalities, scRNA-seq and scTCR-seq, as described above, followed by labeling of T-cell clonal identity for each T cell. The row-normalized, log-transformed test data was then passed to the model, resulting in a learned low-dimensional representation of the data (Figure 4B). Similar to our training data, we calculated the patient clonal purity score over the test data, and compared its values to those obtained by a standard expression-based preprocessing pipeline, using a paired two-sided Wilcoxon signed-rank test (Figure 4D). We also evaluated the Spearman correlation between the observed and the predicted log clone size obtained by the complementary clone-size prediction model (Figure 4E). Lastly, T-cell states/types were annotated as described above (Figure S6A-B), and the Leiden clustering algorithm was applied to identify T-cell clonal niches, as was similarly performed for our training data (Figure 4F). Following manual inspection of multiple clustering resolutions, including 0.5, 0.75, and 1, the resolution of 1 was used to label clonal niches for downstream analysis. Similarity of the cellular composition between the niches obtained in the test data and those obtained in the training data, was then calculated using total variation distance (TVD, Table S3). More explicitly, for each niche we calculated the fractions of the different cell types composing that niche, and used these fractions to calculate the TVD between each niche *p* in the training data and each niche *q* in the test data:

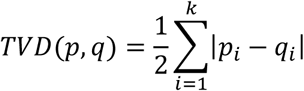

*p_i_* represents the fraction of cell type *i* in niche *p*, *q_i_* represents the fraction of cell type *i* in niche *q*, and *k* represents the total number of cell types, such that lower TVD corresponds to a higher similarity of the cellular composition of both niches.

### Association of clonal niches with patient clinical outcomes

To identify clonal niches that are significantly associated with patient response to therapy, as well as identifying their dynamics following treatment across responding and non-responding patients, we focused on 466 tumor samples obtained from 237 patients, representing 17 different studies of cancer patients treated with immunotherapy across the training data (Table S1). Then, for every clonal niche we calculated the fraction of cells assigned to that niche in every sample and calculated Cliff’s delta, together with its corresponding consistent sampling variance^76^, to compare these fractions between responders and non-responders. The pooled effect estimate for each clonal niche was then calculated using the DerSimonian-Laird method^74^, followed by correction for multiple hypothesis across niches using the Benjamini-Hochberg false discovery rate^93^, considering adjusted p-values < 0.05 as significant (Figures 5F, S7C, Table S4). The DerSimonian-Laird method was applied using only studies containing at least two responders and two non-responders, resulting in 13 eligible studies out of the overall 17 studies with available response annotations. In addition, whenever a specific niche was not present at all across all the samples of a specific study, that study was excluded from the analysis of that specific niche. We performed a follow-up analysis using the clonal niche that was significantly associated with patient response across studies (niche #5), by comparing its fractions between responders and non-responders for all the 17 studies together, using a two-sided Wilcoxon rank-sum test (Figure 5G). This included additional separate comparisons using samples obtained either before or after treatment (Figure 5G).

Among all 237 patients having response annotations, 67 responders and 32 non-responders had longitudinal tumor samples obtained both prior to- and following treatment. We used their samples to test for differences in niche abundance between the two time-points. Whenever a patient had multiple biopsies taken either before or following treatment, we averaged niche abundance across all the samples within that time-point to obtain a single value per time-point. Notably, two NSCLC patients from Liu et al.^13^ had multiple biopsies taken following therapy from different sites having different response statuses. In these cases, we considered the baseline/post-treatment sample pairs that were taken from the same biopsy site and had the same response status. We performed a paired two-sided Wilcoxon signed-rank test to compare the abundance of niche #5 between the two time-points between responders and non-responders (Figure 5H). We applied similar process to patients having both tumor and blood samples, and compared niche #5 abundance between tumor and blood samples for responders and non-responders, separately (Figure 5I). The DerSimonian-Laired meta-analysis was repeated across the training data also at the level of differentiation states, regardless of clonal niche identity (Figure S7D-E, Table S4). We also repeated this entire analysis separately, using the test data, containing four different studies of cancer patients having response annotations (Figure 5L, Table S4).

### Analysis of HBV-dextramer-barcoded single-cell sequencing data

Out of the 11 studies in our test data, we focused on the dataset by Guo et al.^30^, containing paired scRNA/TCR-seq of HBV^+^ HCC patients receiving neoadjuvant anti-PD1 plus Lenvatinib, including HBV-dextramer-barcoded single-cell sequencing information for five patients, which was analyzed using a pre-processing approach similar to the authors’ original pipeline (Figure 6A, Table S1). More specifically, we identified high-confident dextramer binding events using the dextramer-count cutoffs provided by the authors, separating epitope-specific bindings from random bindings. Following, among the confident dextramer bindings per cell, the dominant dextramer per cell was labeled based on the highest dextramer count in that cell. In cases where the same highest count was observed for more than one dextramer per cell, the dextramer with the highest dominant abundance in that cell’s clonotype was chosen. Next, we focused on clones having at least 2 cells with dextramer data and constructed a 2x2 table for each dextramer in each clone. For each clone, this table counted for each dextramer how many cells from that clone dominantly bind that dextramer, how many cells from that clone do not dominantly bind it, as well as how many cells from all other clones dominantly bind or do not bind that dextramer. Fisher’s exact test was then performed on each 2x2 table to calculate the p-value, the odds ratio, and the binding-weighted odds ratio, which was calculated by multiplying the odds ratio by the fraction of the binding cells in the tested clone. Whenever the odds ratio for a given clone-dextramer pair was infinite, it was assigned the highest finite odds ratio observed for that clone, provided that at least one finite positive value was available for that clone. P-values across all tables were then corrected for multiple hypothesis using the Benjamini-Hochberg false discovery rate^93^, further considering clone-dextramer pairs with corrected p-value < 0.05 as significantly enriched. We then extracted all the HBV-specific clones that contain at least one significantly enriched dextramer and calculated a threshold for the binding-weighted odds ratios across all dextramers for these clones, using the 90^th^ percentile. Finally, clone-dextramer pairs were defined as ‘high avidity pairs’ when their corrected p-value was below 0.05 and their binding-weighted odds ratio was above the threshold (Figure 6B). Whenever multiple clone-dextramer pairs were considered as ‘high avidity pairs’ in the same clone, the pair with the highest weighted odds ratio was selected. Importantly, for simplicity and visualization purposes, whenever a clone-dextramer pair was classified as a ‘high avidity pair’, we labeled all the T cells within that clone as binding the corresponding dextramer, although not all cells within the clone were confirmed to bind it, and some may instead bind other dextramers. Similarly, whenever a clone-dextramer pair was not classified as a ‘high avidity pair’, we labeled all the T cells within that clone as not binding the corresponding dextramer, although some specific individual T cells within that clone may bind it.

Following, the abundance of HBV-specific T cells from high-avidity clones was evaluated across clonal niches (Figure 6E). Of note, several samples in this dataset originally included multiple sub-samples that were technically sorted and sequenced separately. In such cases, we merged these technical sub-samples according to their original biological sample identity.

### Analysis of additional datasets with labeling of antigen-specificity

We utilized three additional public scRNA/TCR-seq datasets, including the HNSCC dataset by Eberhardt et al.^32^, the MCC dataset by Pulliam et al.^33^, and the AML dataset by Kim et al.^34^ (Table S1). Importantly, all of these datasets were completely external to model, and were not used in either training or test sets. The HNSCC dataset included only HPV-specific T cells that were sorted prior to sequencing, and its scRNA/TCR-seq data was pre-processed as described above, retaining cells having both scRNA-seq and scTCR-seq that passed our QC. The scRNA/TCR-seq data of the MCC cohort was also preprocessed as described above, with cell-hashing demultiplexing performed using HashSolo^94^. TCRs from cells passing our QC were then mapped to their corresponding antigens using the antigen-specificity labels provided by the authors^33^.

In the AML dataset by Kim et al.^34^, the authors used a hashtag antibody to label dextramer^-^ cells prior to sequencing. For each sample having cell-hashing, we plotted the density curve of the hashtag oligonucleotide counts and defined a per-sample threshold for hash-positive cells as the trough of the bimodal density curve. Following, hashed cells were labeled as dextramer^-^ and un-hashed cells as dextramer^+^, as noted by the authors. We focused on the SRSF2-mutant patient cohort and applied DexTRAC (https://github.com/lijxug/DexTRAC)^30^ to the dextramer-count matrix provided by the authors.

More specifically, we used the dextramer-negative population as a background distribution for random bindings associated with each dextramer, and extracted the expression cutoff for each dextramer that accounts for epitope-specific bindings. Further focusing on dextramer^+^ cells, we identified dominant dextramer per cell and labeled high-avidity T-cell clones as described above. Of note, we analyzed this specific dataset at the level of each technical sample, as no public mapping to patient identities was provided, with scTCR-seq provided only with the CDR3 amino-acid sequences.

CloneTrast was applied to each of the three datasets, followed by projection of the obtained embedding upon the UMAP of the test dataset, including visualization of antigen-specific T cells in each dataset (Figure 6H-J). We then mapped each cell from the three datasets to clonal niches, based on their 15 nearest neighbors from the test data in the CloneTrast embedding space, using the cosine metric (Figure S8A-D).

Finally, we used the NSCLC dataset of Caushi et al.^31^ and labeled antigen-specific T-cell clones based on the labels provided by the authors. As this dataset existed in our training data, we colored the low-dimensional representation obtained using CloneTrast by antigen-specific T cells from this dataset (Figure S8E), and showed their membership across the 18 clonal niches from the training data (Figure S8F).

### Quantification and statistical analysis

The quantitative and statistical analyses are described thoroughly in the relevant sections of the method details. Explicit p-values and sample sizes are embedded in the figures and/or figure legends, and are described throughout the text.

## Supplemental information titles and legends Document S1. Figures S1-S8

**Table S1. Summary of utilized paired scRNA/TCR-seq datasets,** Related to Figures 1-2, 4-6.

**Table S2. Train data information,** Related to Figures 2-3, S2-3.

**Table S3. Test data information,** Related to Figures 4, S6.

**Table S4. Response meta-analysis results,** Related to Figures 5, S7.

